## Supplementary Information for "Engineering *Pseudomonas putida* KT2440 for chain length tailored free fatty acid and oleochemical production"

**Authors & Affiliations**

Luis E. Valencia [1,2,3], Matthew R. Incha[1,2,4], Matthias Schmidt[1,2,5], Allison N. Pearson[1,2,4], Mitchell G. Thompson[1,6], Jacob Roberts[1,2,3], Marina Mehling[1,2], Kevin Yin[1,2,4], Ning Sun[2,7], Asun Oka[2,7], Patrick M. Shih[1,2,4,6], Lars M. Blank[5], John Gladden[1,8], Jay D. Keasling[1,2,3,9-11*]

[1] Joint BioEnergy Institute, Emeryville, CA 94608, USA
[2] Biological Systems and Engineering Division, Lawrence Berkeley National Laboratory, Berkeley, CA 94720, USA
[3] Department of Bioengineering, University of California, Berkeley, CA 94720, USA
[4] Department of Plant and Microbial Biology, University of California, Berkeley, CA 94720, USA
[5] Institute of Applied Microbiology (iAMB), Aachen Biology and Biotechnology (ABBt), RWTH Aachen University, Aachen, Germany
[6] Environmental Genomics and Systems Biology Division, Lawrence Berkeley National Laboratory, Berkeley, California, USA
[7] Advanced Biofuels and Bioproducts Process Demonstration Unit, 5885 Hollis Street, Emeryville, California 94608, USA
[8] Biomanufacturing and Biomaterials Department, Sandia National Laboratories, Livermore, CA 94550, USA
[9] Department of Chemical & Biomolecular Engineering, University of California, Berkeley, CA 94720, USA
[10] Center for Biosustainability, Danish Technological University, Lyngby, Denmark
[11] Center for Synthetic Biochemistry, Institute of Synthetic Biology, Shenzhen Institutes of Advanced Technologies, Shenzhen, China

**Table of Contents**

1. **Figure S1 - CoA Ligase Mutant RB-TnSeq Fitness Data**
2. **Figure S2 - Isovalerate production**
3. **Figure S3 - Carbon source growth assays**
4. **Table S1 - FFA and FAME titers in *P. putida* after growth in 250 mL shake flask cultures**
5. **Table S2 - Strains used in this study**
6. **Table S3 - Plasmids used in this study**

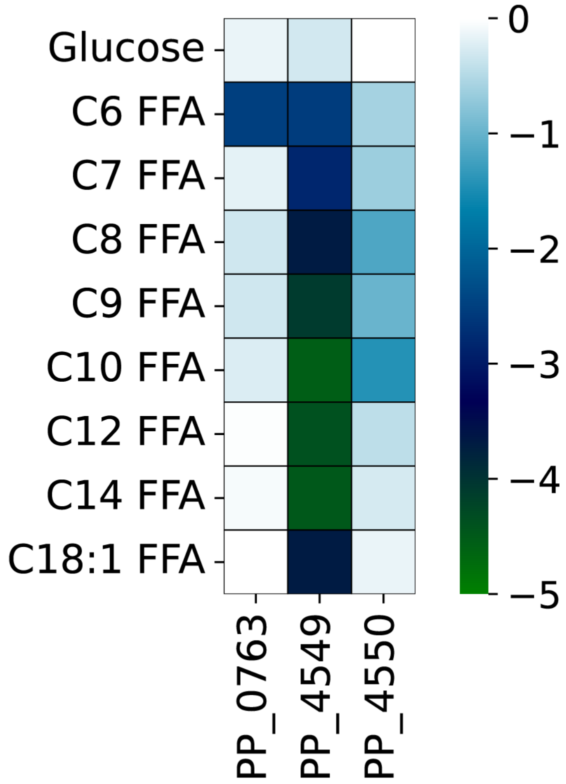


**Figure S1. CoA Ligase Mutant RB-TnSeq Fitness Data.** Heat map showing fitness scores for various CoA ligase knockouts grown minimal media containing free fatty acids or glucose as a sole carbon source. The colors represent fitness scores, with white representing neutral fitness and green representing negative fitness.[1]

A)

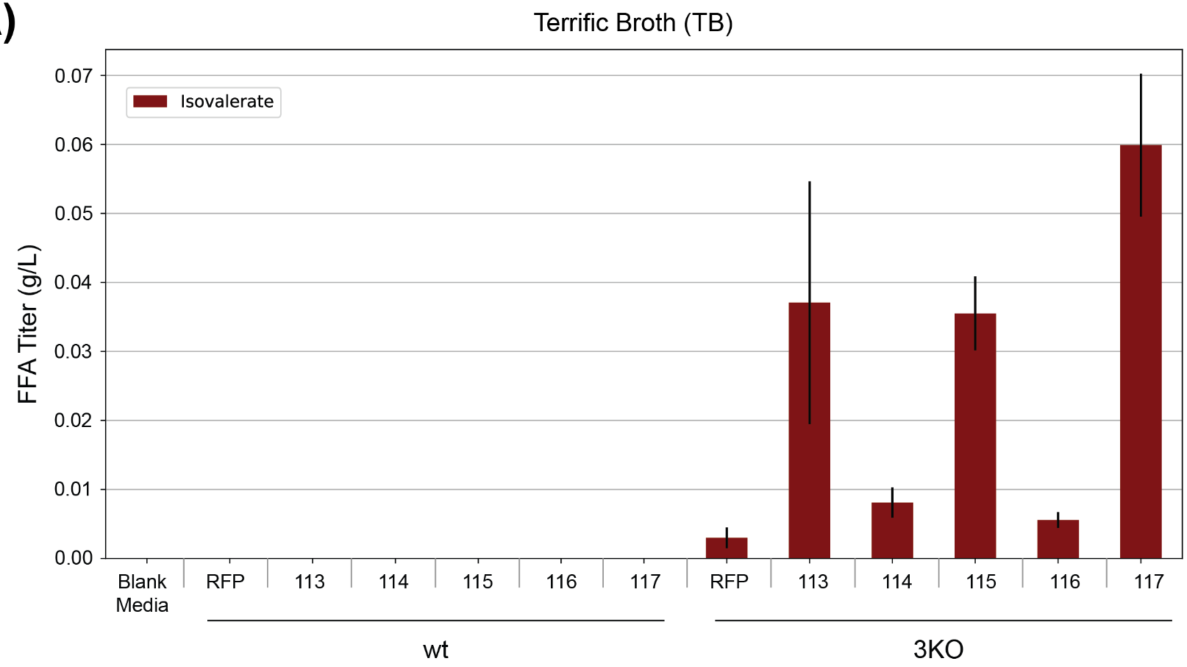


B)

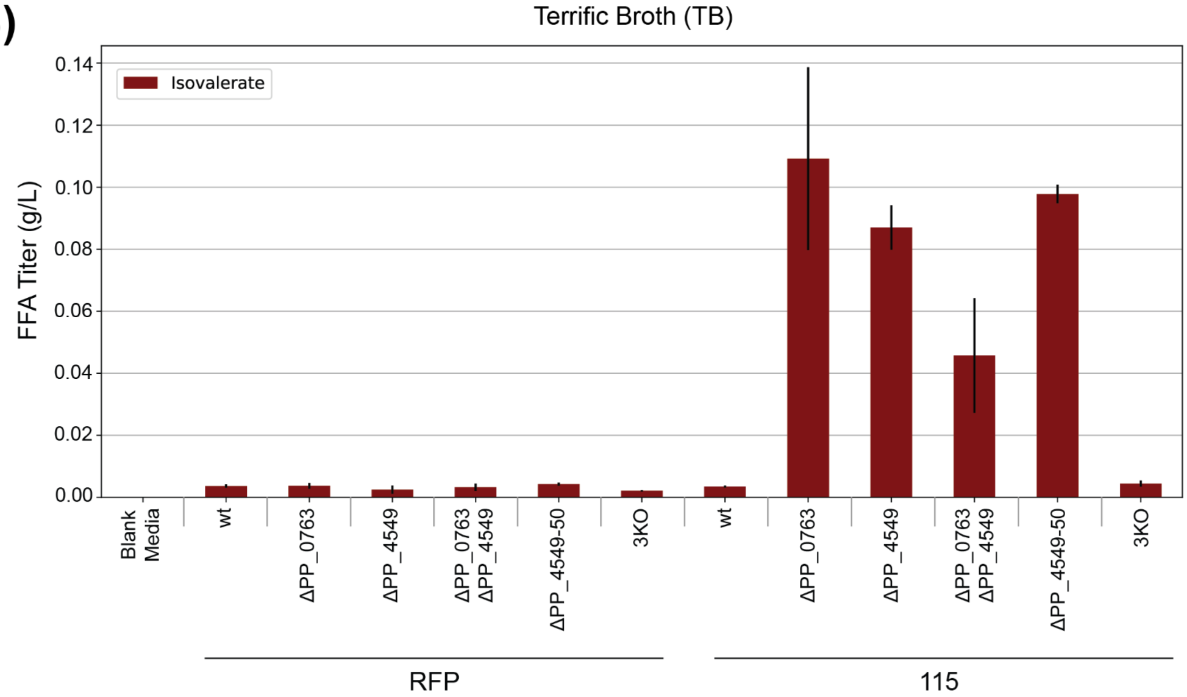


**Figure S2. Isovalerate production in various *P. putida* strains. a** Isovalerate production in Terrific Broth across various ‘TesA variants in wt and 3KO background strains. **b** Isovalerate production across various *P. putida* background strains after overexpression of RFP or ‘TesA R3.M4.

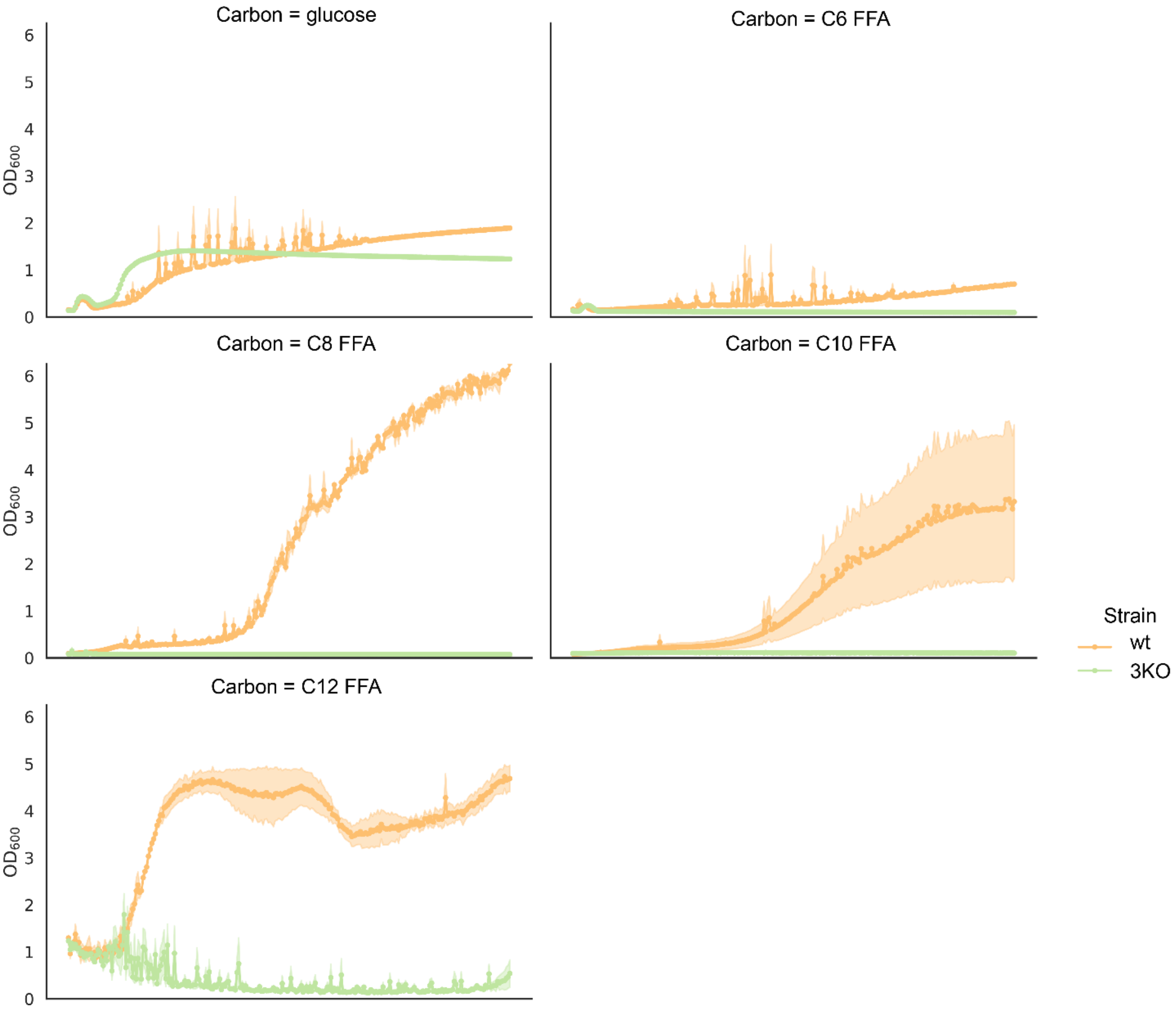


**Figure S3. Growth curves for wt and 3KO on glucose or medium-chain FFAs.**

A)

| Media | Strain | FFA Titer (mg/L) | | | | | | | | | | Titer Fraction Medium-Chain |
|---|---|---|---|---|---|---|---|---|---|---|---|---|
| | | C6 | C8 | C10 | C12:1 | C12 | C14:1 | C14 | C16:1 | C16 | Total | |
| EZ Rich + 100mM Glycerol | Δ0763 Δ4549 + 115 | 21.0 ± 0.1 | 81.5 ± 0.5 | 5.3 ± 0.1 | 21.2 ± 0.2 | 21.7 ± 0.5 | 9.3 ± 0.4 | 1.2 ± 0.4 | 25.4 ± 0.6 | 2.7 ± 0.4 | 189.3 ± 3.0 | 0.80 |
| | 3KO + 115 | 11.9 ± 0.8 | 85.4 ± 5.1 | 5.0 ± 0.5 | 22.7 ± 1.6 | 27.3 ± 2.4 | 9.7 ± 1.2 | 14.5 ± 2.1 | 48.2 ± 1.1 | 34.3 ± 5.0 | 258.9 ± 19.8 | 0.59 |
| | 3KO + RFP | 0.0 ± 0.0 | 0.0 ± 0.0 | 0.0 ± 0.0 | 0.0 ± 0.0 | 0.0 ± 0.0 | 0.0 ± 0.0 | 0.0 ± 0.0 | 33.5 ± 0.3 | 16.0 ± 2.6 | 49.5 ± 2.9 | 0.00 |
| Terrific Broth | Δ0763 Δ4549 + 115 | 47.3 ± 5.2 | 122.3 ± 8.0 | 8.3 ± 1.1 | 19.1 ± 10.4 | 0.0 ± 0.0 | 0.0 ± 0.0 | 0.0 ± 0.0 | 0.0 ± 0.0 | 0.0 ± 0.0 | 197.0 ± 24.7 | 1.00 |
| | 3KO + 115 | 36.3 ± 5.1 | 253.6 ± 3.1 | 26.4 ± 0.9 | 84.3 ± 4.0 | 97.5 ± 6.6 | 30.3 ± 1.3 | 53.9 ± 3.3 | 50.1 ± 4.6 | 38.4 ± 2.3 | 670.9 ± 31.3 | 0.74 |
| | 3KO + RFP | 0.0 ± 0.0 | 0.6 ± 0.0 | 0.0 ± 0.0 | 0.0 ± 0.0 | 0.7 ± 0.2 | 5.9 ± 1.0 | 11.8 ± 1.4 | 183.1 ± 1.8 | 395.1 ± 24.0 | 597.2 ± 28.4 | 0.00 |
| Diluted Dry Sorghum Hydrolysate | Δ0763 Δ4549 + 115 | 43.0 ± 1.4 | 210.2 ± 4.0 | 26.2 ± 0.9 | 87.7 ± 1.7 | 79.8 ± 4.0 | 13.3 ± 4.7 | 46.6 ± 10.4 | 19.5 ± 4.6 | 35.0 ± 4.8 | 561.1 ± 36.5 | 0.80 |
| | 3KO + 115 | 1.4 ± 0.8 | 147.9 ± 1.7 | 13.7 ± 0.6 | 54.5 ± 0.8 | 75.4 ± 1.3 | 31.5 ± 0.4 | 60.7 ± 3.2 | 49.5 ± 3.6 | 51.5 ± 2.6 | 486.1 ± 15.0 | 0.60 |
| | 3KO + RFP | 0.0 ± 0.0 | 10.4 ± 3.5 | 0.0 ± 0.0 | 0.0 ± 0.0 | 3.0 ± 0.5 | 3.6 ± 0.3 | 5.4 ± 0.5 | 16.7 ± 1.1 | 47.8 ± 2.3 | 86.9 ± 8.2 | 0.15 |

B)

| Media | Strain | FAME Titer (mg/L) | | | | | |
|---|---|---|---|---|---|---|---|
| | | C6 | C8 | C10 | C12:1 | C12 | Total |
| EZ Rich + 100mM Glycerol | Δ0763 Δ4549 + 125 | 1.0 ± 0.4 | 23.1 ± 3.4 | 0.8 ± 0.4 | 5.8 ± 0.3 | 0.0 ± 0.0 | 30.7 ± 4.5 |
| | 3KO + 125 | 1.7 ± 0.4 | 23.5 ± 5.9 | 0.0 ± 0.0 | 3.4 ± 0.5 | 0.0 ± 0.0 | 28.6 ± 6.8 |
| Terrific Broth | Δ0763 Δ4549 + 125 | 25.8 ± 0.3 | 199.0 ± 10.7 | 22.4 ± 1.3 | 35.0 ± 0.9 | 20.1 ± 1.1 | 302.4 ± 14.2 |
| | 3KO + 125 | 25.0 ± 0.9 | 187.0 ± 13.0 | 15.6 ± 2.0 | 30.6 ± 2.4 | 20.3 ± 2.0 | 278.5 ± 20.2 |
| Diluted Dry Sorghum Hydrolysate | Δ0763 Δ4549 + 125 | 11.7 ± 1.3 | 43.0 ± 6.0 | 2.1 ± 1.4 | 5.7 ± 0.8 | 1.0 ± 1.0 | 63.4 ± 10.5 |
| | 3KO + 125 | 27.0 ± 1.6 | 167.1 ± 10.8 | 8.9 ± 1.6 | 12.1 ± 0.8 | 8.9 ± 1.4 | 224.0 ± 16.1 |

**Table S1. FFA and FAME titers in various *P. putida* strains after growth in 250 mL shake flask cultures. a** FFA titers after growth in 250 mL shake flasks. The fraction of the total titer that was composed of medium-chain FFAs is reported in the final column. **b** FAME titers after growth in 250 mL shake flasks. Values shown represent the mean of biologically independent samples and the standard error of the mean (n = 3).

| Strain | Description | JBEI part ID | Reference |
|---|---|---|---|
| *E. coli* XL1 Blue | | | Agilent |
| *E. coli* S17-1 λpir | | | [2] |
| *P. putida* KT2440 | Wildtype | | ATCC 47054 |
| *P. putida* ΔPP_0763 | Strain with complete internal in-frame deletion of PP_0763 | JPUB_020583 | This study |
| *P. putida* ΔPP_4549 | Strain with complete internal in-frame deletion of PP_4549 | JPUB_020584 | This study |
| *P. putida* ΔPP_0763 ΔPP_4549 | Double knockout strain with complete internal in-frame deletion of PP_0763 and PP_4549 | JPUB_020585 | This study |
| *P. putida* ΔPP_4549-50 | Double knockout strain with complete internal in-frame deletion of PP_4549 and PP_4550 | JPUB_020586 | This study |
| *P. putida* ΔPP_0763 ΔPP_4549-50 | Triple knockout strain with complete internal in-frame deletion of PP_0763, PP_4549, and PP_4550. Alias 3KO. | JPUB_020587 | This study |

**Table S2. Strains used in this study.**

| Plasmid | Description | JBEI part ID | Reference |
|---|---|---|---|
| pBADT | Broad host-range arabinose inducible plasmid | | [3] |
| pBADT-RFP | pBADT derivative harboring RFP | | [3] |
| 113 | pBADT derivative harboring wildtype ‘TesA | JPUB_020588 | This study |
| 114 | pBADT derivative harboring ‘TesA variant L109P | JPUB_020589 | This study |
| 115 | pBADT derivative harboring ‘TesA variant R3.M4 | JPUB_020590 | This study |
| 116 | pBADT derivative harboring ‘TesA variant CM-5 | JPUB_020591 | This study |
| 117 | pBADT derivative harboring ‘TesA variant RD-2 | JPUB_020592 | This study |
| 125 | pBADT derivative harboring ‘TesA variant R3.M4 and MmFAMT, a fatty acid methyltransferase from *Mycobacterium marinum* | JPUB_020593 | This study |
| pMQ30 | Suicide vector for allelic replacement of $Gm^r$, SacB | | [4] |
| pMQ30 ΔPP_0763 | pMQ30 derivative harboring 1-kb flanking regions of PP_0763 | JPUB_020580 | This study |
| pMQ30 ΔPP_4549 | pMQ30 derivative harboring 1-kb flanking regions of PP_4549 | JPUB_020582 | This study |
| pMQ30 ΔPP_4549-50 | pMQ30 derivative harboring 1-kb flanking regions of PP_4549 and PP_4550 | JPUB_020581 | This study |

**Table S3. Plasmids used in this study.**
